# Host genetics and social relationships jointly shape fitness-associated microbiome variation in a population of feral horses

**DOI:** 10.64898/2026.07.27.740960

**Authors:** Mason R. Stothart, Philip D. McLoughlin, Alastair J. Wilson, Jocelyn Poissant

## Abstract

Gut microbiome variation is consequential for host fitness, but its contribution to adaptive evolution in host populations remains unclear. Although gut microbiome heritability has been demonstrated in humans, domesticated animals, and wildlife, no study has simultaneously assessed heritability of, and natural selection on, microbiome traits. Moreover, prior studies have not investigated whether non-genetic inheritance mechanisms (e.g., maternal and social transmission) could also contribute to microbiome-mediated adaptation. Using quantitative genetic animal models and shotgun metagenomic data from the long-term, individual-based study of feral Sable Island horses (2394 samples, 794 known-fate individuals), we estimated additive genetic, maternal, and social contributions to fitness-associated dimensions of the microbiome identified using canonical analyses of principal coordinates. Combined, additive genetic, permanent environment, and social effects explained 47% and 38% of the variance in fitness-associated dimensions of microbiota or microbial gene family profiles, respectively, with social effects being 2−4 times stronger than additive genetic effects on average. Contrary to our predictions, we observed no evidence for maternal effects. We provide the first evidence that fitness-associated microbiome traits are heritable, and so, theoretically capable of evolution in response to selection. Strong social effects further indicate the possible concurrent importance of non-genetic inheritance to microbiome-mediated adaptive evolution.

## Introduction

Gut microbiomes are hypothesized to underlie phenotypic and fitness variation within animal populations and therefore facilitate adaptation^1–3^. The existence of microbiome-mediated adaptation in nature is supported by theory^3–5^, laboratory experiments^6–8^, the apparent importance of gut microbiomes to the ecology of their host (e.g., herbivory^9^, diet detoxification^10^, hibernation^11^), and patterns of phylosymbiosis at macroevolutionary scales^12^. However, relationships between host fitness and microbiome traits and the mechanisms underlying microbiome trait inheritance (heritability, social transmissibility) have yet to be simultaneously characterized in a wild animal population.

Studies of wild animal populations have recently reported evidence of selection acting on microbiome traits (e.g., alpha diversity, community principal components, microorganism taxon or microbial gene abundance). Hypothesized mechanisms for observed relationships between the gut microbiome and host survival (a fitness proxy) include immunomodulation and energy harvest^13^, pathogen defense^14^, and pathogenesis or loss of digestible energy to methane emissions^15^. However, because the heritabilities of fitness-related microbiome traits are unknown, so too are their capacities to evolve in response to selection.

Quantitative genetic studies of microbiome heritability provide complementary evidence to research on microbiome-fitness relationships, and have yielded similar findings across human, domestic animal, and wildlife populations^16^. While microbiome traits tend to be weakly heritable on average^17,18^, in some cases they are as strongly heritable as conventional animal traits. Yet, it is unclear what reported patterns of microbiome heritability mean for adaptive host-microbiome evolution in nature because studies of microbiome heritability have lacked paired estimates of selection. Specifically, the average heritability of microbiome traits may be less important to understanding microbiome-mediated adaptive evolution than the heritability of microbiome traits that are under selection. Moreover, while previous studies have focused on narrow-sense heritability to infer evolutionary potential, this may not be sufficient to predict intergenerational change for traits also influenced by maternal and/or social transmission. This is because, if present, non-genetic mechanisms may also contribute to microbiome inheritance^19,20^.

Host-to-host transmission of microbes through social networks is now emerging as a critical determinant of microbiome community structure within animal populations^21,22^, and likely contributes to the intergenerational transmission of microbiome variation. For example, in humans, co-habitation and social intimacy creates signatures of bacterial strain-sharing that can persist for decades after separation^23,24^. Similar social effects are observed among both group-living^25–27^ and non-group-living wild mammals^28,29^. Microbe dispersal between animals can occur if they overlap in time and space, and transmission can be further reinforced through social behaviours (grooming, nursing, coprophagy, antagonistic interactions, mating) which can transmit different microbes than those acquired from the environment^30^.

Gut microbiome traits have been likened to a socially transmissible phenotype^31^, but one whose inheritance is shaped by metacommunity ecology dynamics^32^. Microbe dispersal through microbial metacommunities might therefore be viewed both as a non-genetic inheritance mechanism, and as a confounder of heritability estimates, if for example, genetic and social relationships covary within populations^21^. Accounting for social transmission of microbes is therefore important for understanding the inheritance of microbiome traits in wild animals^20^. But what measure of microbiome trait variation is most salient to understanding host-microbiome evolution?

Some hypotheses of host-microbiome evolution suggest emphasis should be placed on microbiome function rather than microbiota community structure. For example, the ‘It’s-the-song-not-the-singer’ [ITSNTS] hypothesis posits that host fitness should be more directly linked to metabolic properties of the microbiome than microbial community membership^33^. Similarly, the ‘ecosystem on a leash’ hypothesis proposes that host physiology is mostly blind to microbe identity when shaping the microbiome, with hosts instead regulating communities by promoting or supressing certain metabolic activities^34^. Despite acting to promote or suppress the same metabolic activities, different hosts within a population may arrive at the same functional outcome via microbiota communities that differ in composition due to functional redundancy in the microbiome^35^. Widespread functional redundancy in the microbiome suggests that measures of microbiota community structure are more sensitive to microbe dispersal limitation and stochastic ecological drift than measures of microbiome function^36^. Neutral ecological processes (drift and dispersal) are non-genetic sources of phenotypic variation which may more strongly reduce the heritability of microbiome traits that reflect microbiota community structure than function^37^. Conversely, measures of microbiome function should be more proximately affected by selective pressures exerted on microbe populations by heritable dimensions of host physiology^34^. Therefore, measures of microbiome function may be simultaneously more strongly heritable (ecosystem-on-a-leash) and more directly connected to host fitness outcomes (ITSNTS), compared to measures of microbiota community structure. Nonetheless, studies of the causes and consequences of gut microbiome variation in the wild have tended to focus on measures of microbiota community structure.

Few free-living animal populations are tractable for the detailed longitudinal monitoring and microbiome sampling required to robustly test hypotheses about microbiome contributions to animal adaptation. The long-term individual-based study of Sable Island feral horses (*Equus caballus*) provides a remarkable opportunity to simultaneously assess the causes and fitness consequences of microbiome variation in the wild^15^. Every horse on Sable Island (∼450-550 horses annually) is monitored from birth to death (known reproductive and survival outcomes without resighting error since 2011) alongside the opportunistic collection of fresh individual-linked fecal samples. The maintenance of a population pedigree, and the segregation of horses into discrete social groups with overlapping home ranges, provides the uncommon opportunity to separate maternal, shared environment, social, and additive genetic contributions to fitness-related gut microbiome traits. As obligate hindgut fermenters, we expect fecal samples to reflect the microbiota that horses rely on to digest and extract nutrients from their fibrous plant-based diet, and therefore fuel survival and reproduction^38^. Indeed, previous analyses have linked fecal microbiome traits to overwinter survival in Sable Island horses^15^.

We sought to quantify the heritability of gut microbiome traits linked to host fitness components (survival and reproductive success) in the long-term individual-based study of Sable Island feral horses (2394 faecal samples, 794 free-living individuals, 7 years of collection), while simultaneously parsing maternal, social, spatial, and shared environment effects using maternity data and observations of social co-occurrence (25,877 horse observation events). By characterizing faecal microbiomes using shallow shotgun metagenomic sequencing^39^, we had the secondary objective to compare selection and quantitative genetic parameter estimates related to traits for microbiota community structure versus microbiome functional potential.

## Methods

### Sable Island Study System

Free-living horses were introduced to Sable Island (Nova Scotia, Canada) in the mid-1700s^40^ and became the subject of a long-term individual-based ecological study in 2007. This long-term study is comprised of an annual census of the feral horse population over ∼2 months, between mid-July and early-September^41,42^. To complete exhaustive annual censuses, the 42-km longitudinal breadth of the island is divided into seven sections of approximately equal size. A minimum of one section is surveyed on foot by researchers per survey day, allowing for complete coverage of the island once per week. During surveys, detailed notes, location (using handheld GPS devices), and photographs are taken for every horse as encountered. Using a combination of colouring markings, scarring patterns, sex, and unique morphological features (e.g., hoof curls, umbilical hernias, chestnut morphology), every individual can be identified through comparisons with a comprehensive photograph database. The population varied between 380–590 horses over the 2013–2020 study period, and 1062 individuals were observed during this time.

Individuals in the Sable Island horse population segregated into 139–171 discrete social groups per year between the 2013–2020 period corresponding to this study, with the number and size of social groups related to an increasingly male-skewed adult sex ratio^43^. Females are invariably found in mixed-sex social groups, in which mating access is usually guarded by a single dominant stallion (mean group size = 5.2 ± 2.3 SD)^44^. Immature males or subordinate males are sometimes observed within mixed-sex groups, but more often, adult males unable to secure harems will remain solitary or form bachelor male social groups (mean group size = 3.4 ± 1.8 SD). The memberships of mixed-sex social groups remain largely unchanged during population censuses, but bachelor male social groups are more dynamic. Social groups do not defend exclusive territories, but rather, occupy discrete overlapping home-ranges. Although environmental differences occur between home-ranges, marram grass (*Calamagrostis breviligulata*) is a core component of the Sable Island horse diet and major differences in diet observed across this population can be described by a quadratic relationship with longitude^15^.

Offspring are typically ejected from mixed-sex social groups at maturity and approximately 30% of females transition among social groups between years, with higher rates of social dispersal at higher population density^44,45^. The inter-annual movement of horses between social groups allows for gene flow and reduces confounding of social and genetic effects. Genetic relationships between horses can be estimated using the population’s social pedigree, constructed using behavioral observations. We use nursing behaviour to confirm maternity and assign paternity to the dominant stallion of a mother’s social group in the year preceding birth, since gestation in horses is 11–12 months in length^46^. An inverse *additive genetic relatedness* matrix and *maternity* data obtained from the population’s pedigree allow us to use quantitative genetic animal models (detailed below) to estimate additive genetic (narrow-sense heritability) and maternal contributions to phenotypic trait variation^47^.

Unobserved extra-pair paternity may increase error in our estimates of genetic relatedness, and therefore, cause a downward bias in estimates of heritability^48^. However, while the rate of extra-pair paternity in Sable Island horses is unknown, such events appear rare in other wild horse populations. For example, an extra-pair paternity rate of ∼2% is observed among feral Camargue horses^49^. The viability of extra-pair offspring in other horses appears limited by infanticide and high maternal costs^50^. Similar patterns of infanticide are observed among Cape mountain zebra (*Equus zebra zebra*), suggesting that female fidelity and extra-pair infanticide may represent an evolved behavioural response in equids^51^.

Fresh faecal samples were collected during population surveys and linked to detailed photos of the sampled individual. All samples were collected within 5 min of defecation using clean nitrile gloves and faeces contaminated with substrate avoided. Faecal samples were kept on ice in the field for a maximum of ∼6 hours, before being subset into microcentrifuge tubes and stored at –20°C on Sable Island for up to 2 months. Samples were placed in long-term storage at –80°C on the mainland after transport from Sable Island^52^. Only samples collected from horses ≥1 year of age and which passed sequencing quality control thresholds were used in analyses, resulting in 2,394 faecal microbiome samples from 794 individuals collected between 2013–2019, with every individual within this dataset represented by a maximum of one sample per year.

### Microbiome Dataset Creation

We analysed a shallow shotgun metagenomic microbiome dataset described in detail by Stothart et al.^15^. In brief, we extracted DNA from ∼0.16 grams of faecal material (2,820 samples, 30 negative controls, 30 ZymoBIOMICS Microbial Community Standard II positive controls) using Qiagen’s QIAamp 96 PowerFecal QIAcube HT kits and manufacturer specifications. Shotgun metagenomic libraries were prepared using iGenomx Riptide high-throughput library preparation kits and sequenced on an Illumina NovaSeq 6000 platform (300 cycles) to an average depth of ∼4 million read pairs per sample.

TRIMMOMATIC and BOWTIE2^53–55^ were used to trim adapters and low-quality bases, remove unpaired reads, and filter quality-controlled reads against the EquCab3 domestic horse reference genome^56^. KAIJU was used to assign microbial taxonomy to quality-controlled read pairs based on translated read hits to the NCBI BLAST non-redundant protein database^57^. Only samples which surpassed 0.4 million assigned read pairs were retained for analysis, as this depth has been validated for the Sable Island feral horse system^39^. Gene family contents were estimated by using HUMANN3 to map translated reads against the UniProt50 protein database^53,58^. Read hits were normalized by gene region size and grouped to MetaCyc gene family reactions for analysis^59^. See Stothart et al.^15^ for greater details about dataset creation.

### Estimates of spatial, shared environment and social relationships

In addition to estimating maternal and additive genetic contributions to fitness-related microbiome traits, we sought to quantify environmental and social effects. Hosts occupying the same environment can converge upon a similar microbiome structure due to similarities in diet, water quality, or local environmental stressors^60^. To quantify spatial autocorrelation in gut microbiome structure, we calculated a *spatial proximity matrix* which describes the geospatial distance separating locations of faecal sample collection (inverse distance with values scaled between 0-1). This matrix was used to estimate the effects of persistent spatial heterogeneity in the environment on horse microbiome trait variation (animal model parameterization detailed below).

As a dynamic dune grass ecosystem, environmental conditions at fixed locations on Sable Island can change across years^61^. Inter-annual differences in horse population densities, climate, or adverse weather conditions might cause environmental differences at the same location between years. Moreover, horses are mobile animals, and so, the *spatial proximity* matrix will not fully reflect shared environments in the time preceding faecal sample collection. To further account for shared environment effects, we therefore assigned each discrete social group of horses a unique *group identity* specific to the year of observation. This *group identity* term is aspatial but delineates cohorts of horses who experienced the same environment during, and in the months preceding, population surveys and faecal sample collection. The memberships of mixed-sex social groups are highly stable within years and easily delineated^44^. Social groups formed by bachelor males are likewise repeatable within years but more dynamic. Bachelor males were therefore assigned to their modal social group. If multiple social assemblages were observed with equal frequency, bachelors were assigned to the group with the largest membership. Finally, if alternative groups were equivalent in both size and observation frequency, bachelor males were grouped to the assemblage whose observations spanned the greatest period.

Microbiome similarities between horses co-occurring in space and time can be due to shared environment effects that result in similar diets or stressors, which in turn exert similar selective pressures on microbial communities (deterministic ecological process); however, social interactions between co-occurring individuals may also provide the opportunity for the direct transmission of microbe between hosts (dispersal process). Sable Island horses engage in a range of behaviours likely to facilitate microbe transmission, including consumption of fresh faeces, social grooming using their mouths, olfactory investigation of faeces, defecating near and into shared water sources, and grazing a landscape littered with faeces. To more precisely parse shared environment versus social transmission contributions to microbiome variation, we therefore constructed a *social community* matrix for testing alongside indices more directly tied to shared environments (*spatial proximity* and *group identity*).

In brief, the *social community* matrix quantifies pairwise similarity in social environment between individuals (within and between years), as well as changes in the social environment of the same individual between years. Specifically, for each horse-year observation (e.g., HorseA_2015, HorseA_2016, HorseB_2016), all other horses observed on the island that year were treated as potential social affiliates and assigned a weight equal to their Simple Ratio Index with the focal horse. SRI values estimate the frequency with which pairs of individuals co-occur and has been shown to correlate with microbiome beta diversity in the wild^29^. SRIs were calculated within each survey year using daily surveys as observation windows. Pairs of horses were considered to co-occur when their recorded locations on the same day were within 300 m, a threshold approximating mean daily movements in this population^26^. For each pair, SRI therefore reflects the proportion of days on which individuals were observed together within a survey year, relative to the total number of days on which one or both were observed. Horses within the same social group are almost always observed within 300 m of one another, except when dominant stallions are briefly absent while engaging rivals. Although this definition does not guarantee direct interactions, it reflects social tolerance and the potential for microbe transmission via indirect pathways such as coprophagy or incidental faecal exposure via forage or water. The table containing year-specific SRI estimates between every pair of horses was used to construct a Euclidean distance matrix that describes pairwise dissimilarities in social relationships. The Euclidean distance matrix was then converted to a social community similarity matrix scaled between 0 and 1 for use in animal model analyses 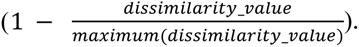

S*patial proximity*, *group identity*, and *social community* estimates each capture a combination of shared environment and social effects, but to differing extents. For example, two horses sampled at the same location in different years may share a similar environment despite never directly interacting and existing in different social environments. Similarly, the *group identity* term delineates horses who tightly co-occur in space and time within a survey year, but does not capture relationships between groups, the more fluid social relationships observed among bachelors, or similarity in group membership between years. In contrast, the *social community* matrix is aspatial and atemporal and more directly estimates similarities in social relationship structure within and between groups, both within and between survey years. As a result, individuals sampled in different years and locations may nonetheless share similar social environments. If social effects on the microbiome are not entirely attributable to shared environment effects—for example, if microbiome variation is transmitted and maintained via microbe dispersal through social networks—then we expect the *social community* matrix to explain variation in the microbiome beyond that captured by *spatial* proximity and *group identity*.

### Statistical Methods

#### Quantifying fitness-related microbiome traits

Previous analysis of the Sable Island horse microbiome identified 155 microbial taxa and 27 gene families significantly associated with overwinter survival^15^. The large number of microbiome traits identified as survival-associated makes previous findings unwieldy for quantitative genetic analysis and interpretation. Furthermore, redundant biological information is likely contained among these results; for example, many of the microbes and gene families identified could contribute to the same pathogenesis and metabolic (methanogenesis) processes hypothesized to connect the gut microbiome to horse survival on Sable Island^15^.

To collapse redundancy in the microbiome patterns we previously detailed into a single ‘survival-associated’ dimension of the gut microbiome, we conducted canonical analyses of principal coordinates (CAPs) using the R package ‘vegan’ and ‘phyloseq’^62,63^. Specifically, we conducted CAPs for microbiota community and gene family composition (Aitchison distances) with a constraint for overwinter survival (individual survived versus died overwinter). CAP analyses were conditioned on DNA extraction plate to control for technical differences between batches of DNA extraction and library preparation. This allowed us to identify axes of microbiome variation strongly associated with horse survival. Weighted average scores from CAP analyses provided us with focal microbiota and microbial gene family measures that comprehensively reflect previous findings^15^ and can be used as focal traits in quantitative genetic analyses. Statistical significance of survival-associated CAP axes was evaluated using analysis of variance tests (ANOVA; aov() parameters: by = “axes”).

Associations between CAP scores and horse survival could be caused by confounding life history or environmental factors. For example, senescent horses may harbour distinct microbiomes, but their increased likelihood of mortality might not be causally connected to the microbiome. We therefore tested whether CAP axes were associated with horse survival independent of life history and environmental factors by using binomial mixed effects models (family=binomial(link=’logit’)) in which overwinter survival [survived / died overwinter] was treated as a response variable, and microbiota or gene family survival-associated CAP scores were treated as predictor variables, alongside year as a random effect, and fixed effects for sex, longitude of sample collection (linear and quadratic terms), age (linear and quadratic terms), and a sex-specific interaction with parental status (foal offspring present during the year of sample collection) to account for the sex-specific energetic requirements for nursing and reproduction. Estimates from this model provide a more comprehensive estimate of the strength of the association between survival and the gut microbiome, when compared to the microbe and gene family specific results reported in Stothart et al.^15^.

Previous analyses^15^, and the CAP analyses detailed above, emphasized survival to the neglect of reproductive success, and yet, evolutionary fitness is shaped by both of these key fitness components. We expected that the hypothesized primary mechanism linking the gut microbiome to survival among Sable Island horses—methane-related loss of digestible energy—should also be consequential for reproductive success. As an independent assessment for the biological significance of survival-linked CAP axes, we therefore tested for relationships between CAP trait measures and horse reproduction using binomial mixed effects models as detailed above. These models included reproductive success in year following sample collection as the response variable (no offspring/offspring produced), fixed effects for sex, longitude of sample collection (linear and quadratic terms), and age (linear and quadratic terms), and a random effect for year. However, unlike survival, we expect the energetic cost of reproduction in the time following sample collection to be higher for females than males, due to the demands of gestation and nursing. Therefore, binomial models for reproductive success included an interaction between sex and CAP trait measures. These tests are not an exhaustive exploration of relationships between the gut microbiome and horse reproductive success. Rather, they are intended to test whether previously detailed patterns, and the foci of this study, are strictly survival-associated, or have a more comprehensive connection to host fitness.

#### Quantitative genetic analysis of microbiome traits

We used quantitative genetic animal models to test for the relative contributions of permanent environment, additive genetics, maternal identity, spatial effects, social group identity, and social community to survival associated CAP traits^37^. Specifically, we treated microbiota and gene family CAP scores as response variables within restricted maximum likelihood linear mixed effects animal models using the R package ASREML-R (version 4.1.0.176). All models contained fixed effects for sex, longitude of sample collection (linear and quadratic), day of year (linear term to account for seasonal variation), year of sample collection (factor), age (linear and quadratic terms to account for microbiome maturation and senescence), and extraction plate (factor). Genetic, shared environment, and social contributions were then quantified by comparing models with differing combinations of random effects (detailed below).

To test for maternal effects, we subset our data to pedigreed individuals with a known maternal identity (1437 samples, 461 individuals) and compared models which contained a random effect for *horse identity* alongside either *additive genetic relatedness* or *maternal identity* terms, against a model containing all three terms. To test for additive genetic, shared environment, and social effects, we expanded our dataset to include samples from all pedigree informative individuals (2013 samples, 892 informative individuals, 650 phenotyped individuals, 622 maternal links, 461 paternal links). We began with a full model which included terms for *horse identity, additive genetics* (A^-1^ matrix), *spatial proximity*, *group identity*, and *social community* among its random effects structure. Proportional contributions of random effects (i.e. intraclass correlations) were estimated by dividing variance components by the phenotypic variance (V_P_), conditional on the fixed effects detailed above. Statistical significance for variance components was determined using likelihood ratio tests between models which did, versus did not, include the focal random effect (1 DF). Resultant p-values were halved to approximate a 50:50 mixture of χ^2^(1 DF) and χ^2^(0 DF), since variance estimates are constrained to ≥ 0^64,65^. Finally, upon identifying the most parsimonious model structure, we quantified the mechanisms that contributed to covariance between microbiota and gene family CAP axes using bivariate animal models. The statistical significance of covariance estimates was assessed using likelihood ratio tests comparing a full model with an unstructured covariance matrix for a given focal term, to a model in which covariance was constrained to zero.

To characterize microbe or gene family specific sensitivities to permanent environment, additive genetic, or social community effects, we separately modelled microbe (1574 taxa) and gene family (1416 gene families) centered log-ratio (CLR) transformed abundances as response variables within quantitative genetic models parameterized as detailed above. CLR transformations were used to normalize read counts due to the compositional nature of our sequencing dataset^66^. Analyses were restricted to features which were present in at least 75% of samples, and because the application of CLR transformations to zero-inflated data could artificially skew variance estimates, samples containing zero counts for a given feature were omitted from analyses of that feature ^15^. We used likelihood ratio tests to determine significance and applied Benjamini-Hochberg correction to p-values to account for a false discovery rate due to multiple testing (FDR)^67^.

GLMMs were used to estimate relationships between microbe and gene family specific variance component estimates and previously reported relationships between these microbiome traits with horse survival (relationship of feature CLR abundance with log odds of horse survival)^15^. for the purpose of testing for non-independence between selection and the transmissibility of microbiome features. Both microbiota and gene family models contained an additional fixed effect for microbe or gene family average relative abundance (log transformed) to account for increased error in the estimation of low abundance microbiome features, and the bias this may introduce into both selection and animal model analyses. The GLMM for microbe abundance measures included an additional random effect for phylum.

## Results

### Identifying survival associated dimensions of the microbiome

Canonical analysis of principal coordinates constrained by overwinter survival resulted in CAP axes which were significantly associated with microbiome beta diversity (CAP_microbiota_: F_1,2363_ = 2.541, p = 0.003; gene family CAP_gene family_: F_1,2363_ = 1.836, p = 0.01), despite only accounting for 1% and 0.1% of variation in microbiota and gene family Aitchison distances, respectively. Microbe- and gene family-specific scores along survival-constrained CAP axes were positively associated with previously estimated relationships between microbes or gene families and horse survival^15^ among Pearson’s product-moment correlation tests (microbiota: *t* = 21.446, *p* = 2.2e^-16^, *r* = 0.48, Figure S1, Appendix 1; gene family: *t* = 20.185, *p* = 2.2e^-16^, *r* = 0.47, Figure S2, Appendix 2). These results support the use of sample scores from survival-constrained CAP analyses of microbiota community and gene family profiles as informative univariate proxies for previously described relationships between the gut microbiome and Sable Island horse survival.

### Selection Analyses

Survival-associated CAP scores from both microbiota and gene family constrained ordinations were positively predictive of survival (community: β = 0.62 ± 0.07, z = 9.227, p < 2e^-16^; gene family: β = 0.86 ± 0.07, z = 11.569, p = 2e^-16^; Figure 1A), even after controlling for age, sex, longitude, and sex-specific effects of reproduction (Table S1 & S2). Female reproductive success in the year following sample collection was likewise positively associated with the CAP scores from microbiota and gene family constrained ordinations among females (CAP_microbiota_: β = 0.34 ± 0.12, z = 4.236, p = 2.3*e*^-5^; CAP_gene family_: β = 0.49 ± 0.08, z = 6.005, p = 1.91*e*^-9^; Figure 1b; Table S3 & S4). Male reproductive success was associated with the microbiota CAP scores (β = 0.21 ± 0.08, z = 2.485, p = 0.01) but not the CAP scores derived from gene family profiles (β = 0.15 ± 0.08, z = 1.816, p = 0.07). Overall, a one standard deviation increase in a microbiota community-derived CAP axis was associated with an 86% increase in the odds of survival, and a 40% and 23% increase in the odds of female and male reproductive success, respectively. An increase of the same magnitude in gene family CAP scores was associated with a 136% increase in the odds of survival and a 63% increase in the odds of female reproductive success. Observed relationships with both survival and reproductive success suggests that the dimension of microbiome variation we identified may have a comprehensive connection to host fitness.

**Figure 1:**
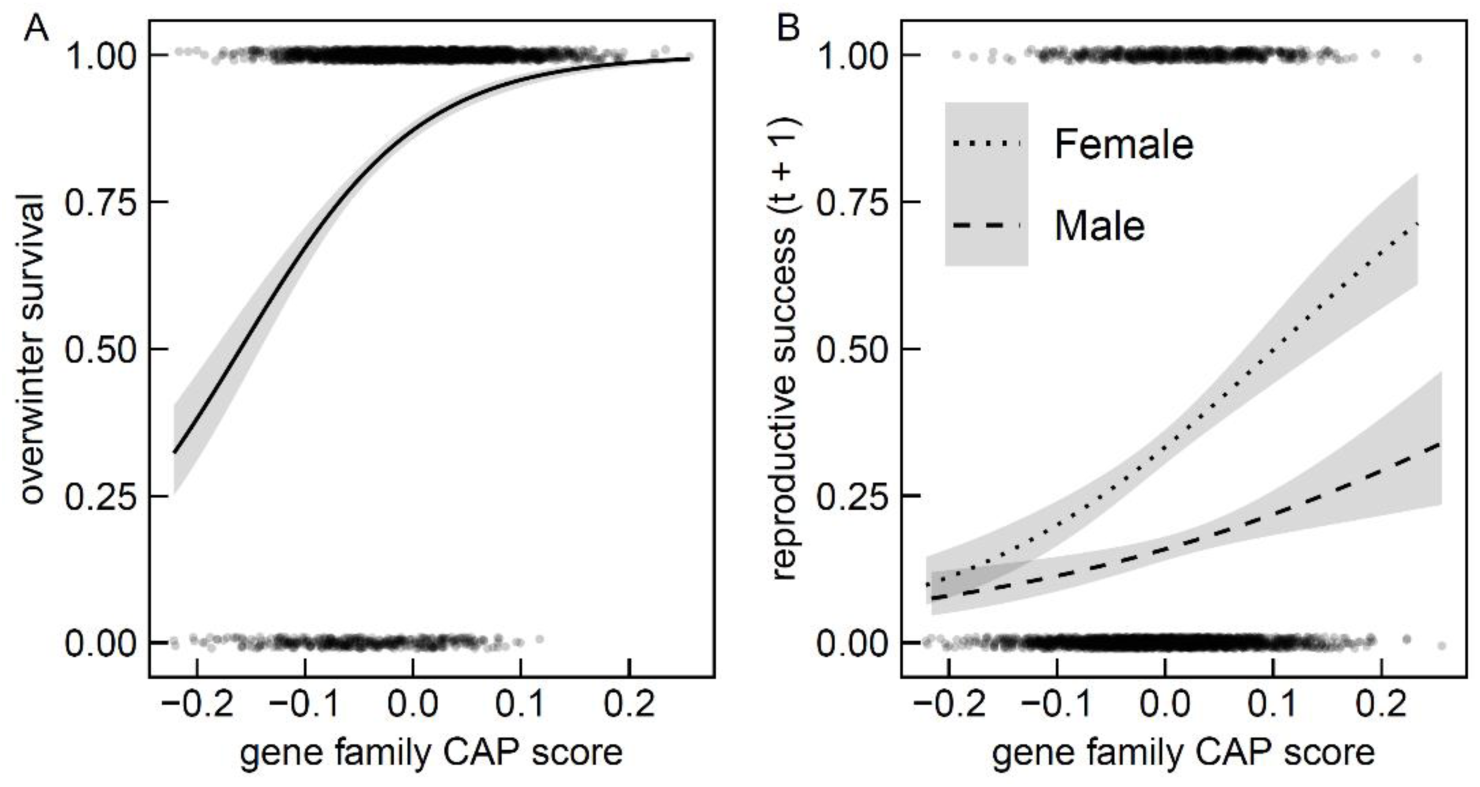
Jittered scatterplots of (a) horse overwinter survival and (b) sex-specific reproductive success in the year following faecal sample collection in response to weighted average scores of samples from a canonical analysis of principal coordinates (CAP) of gene family profiles constrained by overwinter survival. Points denote individual samples with best-fit logistic regression lines and 95% confidence interval shading.

### Quantitative Genetic Analysis

We next sought to partition additive genetic, maternal, spatial, social group, and social community contributions to survival- and reproductive success-associated CAP score variation using quantitative genetic animal models. Amongst a subset of samples collected from horses with a known mother, inclusion of an additive genetic term alongside horse identity and maternal identity significantly improved model performance for the microbiota CAP trait (χ^2^ = 5.20, p = 0.01), but not the CAP scores derived from gene family profiles (χ^2^ = 2.25, p = 0.07). The addition of a random effect for maternal identity to a model containing terms for *horse ID* and *additive genetic* effects did not significantly improve model performance (CAP_microbiota_: χ^2^ ≈ 0, p = 0.5; CAP_gene family_: χ^2^ ≈ 0, p = 0.5). Therefore, we observed some evidence for additive genetic effects, but not maternal identity effects.

Finding no evidence for maternal identity effects, we expanded animal model analyses to include samples from horses of unknown maternity, beginning with a full model that included terms for horse ID, additive genetic, spatial, social group, and social community effects. Likelihood ratio tests provided evidence for permanent environment (χ^2^ = 9.03, *p* = 1.3*e*^-3^), additive genetic (χ^2^ = 7.82, *p* = 2.6*e*^-3^), and social community (χ^2^ = 18.14, *p* = 1.0*e*^-5^) contributions to microbiota CAP score variation, but no evidence for spatial (χ^2^ = 1.15, *p* = 0.14), or social group identity effects (χ^2^ = 0.01, *p* = 0.45; Table 1; Figure 2) when comparing full models to ones in which focal terms were omitted. The most parsimonious model based on AIC analysis included terms for horse ID, additive genetic effects, and social community effects.

**Figure 2:**
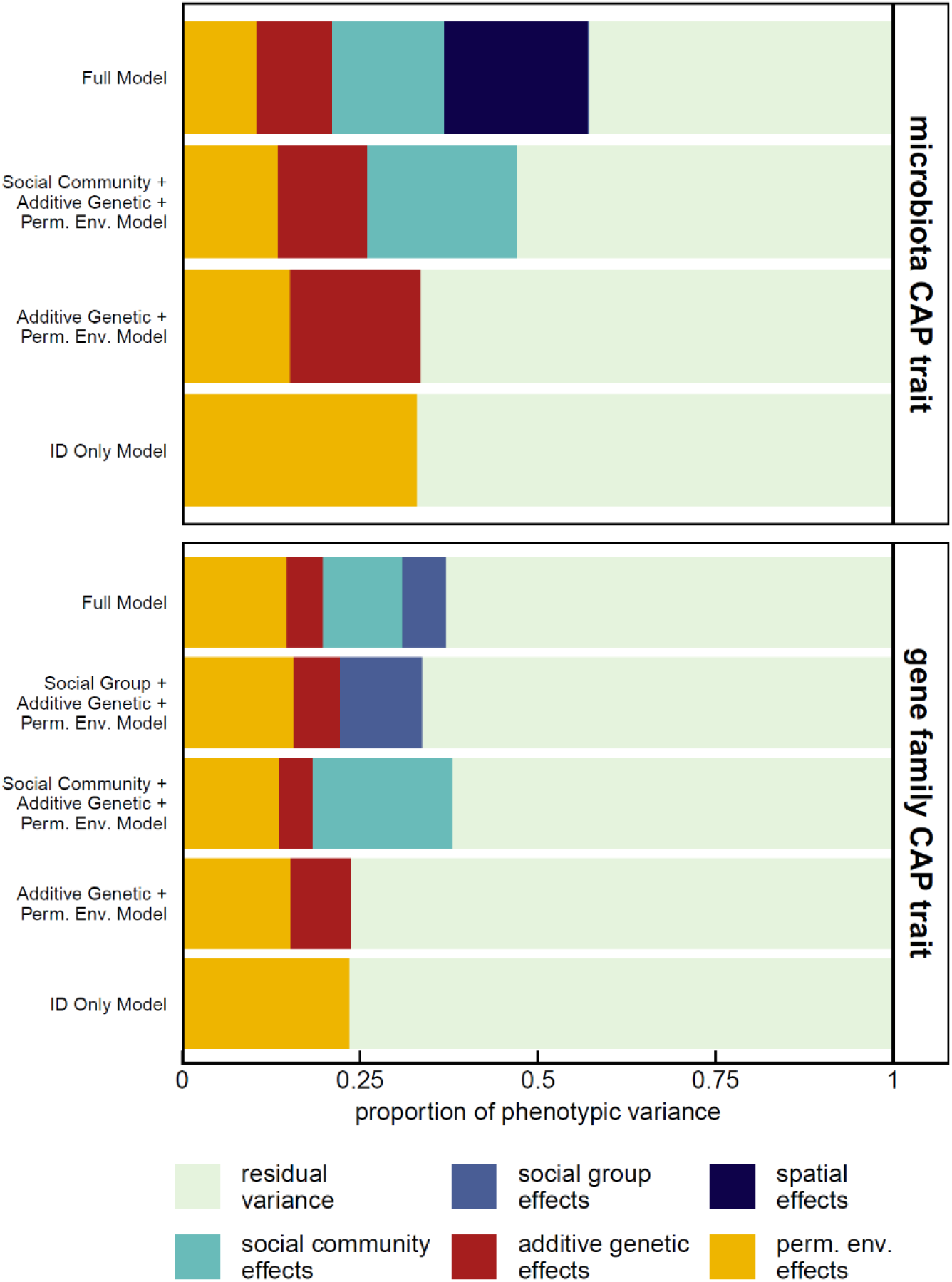
Stacked barplots of proportions of variance in microbiota or gene family derived constrained ordination axes associated with horse overwinter survival that were attributable to additive genetic, permanent environment, spatial, social group identity, or social community effects. Variance component estimates were conditional on fixed effects for longitude (2^nd^ order polynomial), horse age (2^nd^ order polynomial), day of year (2^nd^ order polynomial), DNA extraction/library preparation plate (factor), and year of sample collection (factor).

**Table 1:** Variance component estimates from competing animal models for microbiota CAP score estimates, conditioned on fixed effects for longitude (2nd order polynomial), horse age (2nd order polynomial), day of year (2nd order polynomial), DNA extraction/library preparation plate (factor), and year of sample collection (factor). Raw variance component estimates (V) and proportional contribution towards to phenotypic variance reported with standard error in brackets. Statistically significant estimates are bolded.

| model | V<sub>P</sub> | V<sub>PE</sub> | V<sub>A</sub> | V<sub>SC</sub> | V<sub>SP</sub> | V<sub>SG</sub> | V<sub>R</sub> | pe<sup>2</sup> | h<sup>2</sup> | sc<sup>2</sup> | sp<sup>2</sup> | sg<sup>2</sup> | r<sup>2</sup> | ΔAIC |
| --- | --- | --- | --- | --- | --- | --- | --- | --- | --- | --- | --- | --- | --- | --- |
| horse ID + additive genetic + social community + spatial + social group | 1.08 | **0.11**<br>(0.04) | **0.12**<br>(0.04) | **0.17**<br>(0.05) | 0.22<br>(0.19) | 0.003<br>(0.02) | **0.46**<br>(0.02) | **0.10**<br>(0.04) | **0.11**<br>(0.04) | **0.16**<br>(0.05) | 0.20<br>(0.14) | 0.003<br>(0.02) | **0.43**<br>(0.08) | 2.85 |
| horse ID + additive genetic + social community + spatial | 1.09 | **0.11**<br>(0.04) | **0.12**<br>(0.04) | **0.17**<br>(0.04) | 0.22<br>(0.19) | — | **0.46**<br>(0.02) | **0.10**<br>(0.04) | **0.11**<br>(0.04) | **0.16**<br>(0.04) | 0.20<br>(0.14) | — | **0.43**<br>(0.08) | 0.86 |
| horse ID + additive genetic + social community | 0.88 | **0.12**<br>(0.04) | **0.11**<br>(0.04) | **0.18**<br>(0.04) | — | — | **0.46**<br>(0.02) | **0.13**<br>(0.04) | **0.13**<br>(0.04) | **0.21**<br>(0.04) | — | — | **0.53**<br>(0.03) | 0.00 |
| horse ID + additive genetic | 0.80 | **0.12**<br>(0.04) | **0.15**<br>(0.04) | — | — | — | **0.53**<br>(0.02) | **0.15**<br>(0.05) | **0.19**<br>(0.05) | — | — | — | **0.66**<br>(0.03) | 43.11 |
| horse ID | 0.80 | **0.26**<br>(0.03) | — | — | — | — | **0.53**<br>(0.02) | **0.33**<br>(0.03) | — | — | — | — | **0.67**<br>(0.03) | 52.90 |

PE: permanent environment, A: Additive Genetic, SC: Social Community, SP: Spatial, SG: Social Group

Likelihood ratio tests similarly provided evidence for permanent environment (χ^2^ = 13.26, *p* = 1.4*e*^-4^), social community (χ^2^ = 6.86, *p* = 4.4*e*^-3^), and social group (χ^2^ = 4.28, *p* = 0.02) contributions to gene family CAP traits values, but no evidence for spatial (boundary estimate) or additive genetic (χ^2^ = 1.81, *p* = 0.09) effects when comparing a full model to ones in which focal terms were removed. The top model based on AIC analysis was one that contained both social group and social community effects, alongside horse ID and additive genetic terms, however, this model performed only marginally better than a simpler model containing terms for horse ID, additive genetic effects, social community (△AIC = 2.02; Table 2).

**Table 2:** Variance component estimates from competing animal models for gene family CAP score estimates, conditioned on fixed effects for longitude (2nd order polynomial), horse age (2nd order polynomial), day of year (2nd order polynomial), DNA extraction/library preparation plate (factor), and year of sample collection (factor). Raw variance component estimates and proportional contribution towards to phenotypic variance reported with standard error in brackets. Statistically significant estimates are bolded and ‘boundary’ estimates denoted by ‘*B*’.

| model | V<sub>P</sub> | V<sub>PE</sub> | V<sub>A</sub> | V<sub>SC</sub> | V<sub>SP</sub> | V<sub>SG</sub> | V<sub>R</sub> | pe<sup>2</sup> | h<sup>2</sup> | sc<sup>2</sup> | sp<sup>2</sup> | sg<sup>2</sup> | r<sup>2</sup> | ΔAIC |
| --- | --- | --- | --- | --- | --- | --- | --- | --- | --- | --- | --- | --- | --- | --- |
| horse ID + additive genetic + social community + spatial + social group | 0.84 | **0.12**<br>**(0.03)** | 0.04<br>(0.03) | **0.09**<br>**(0.05)** | *B* | **0.05**<br>**(0.03)** | 0.54<br>(0.03) | **0.15**<br>**(0.04)** | 0.05<br>(0.04) | **0.11**<br>**(0.05)** | *B* | **0.06**<br>**(0.03)** | **0.62**<br>**(0.04)** | 2.00 |
| horse ID + additive genetic + social community + social group | 0.85 | **0.12**<br>**(0.03)** | 0.04<br>(0.03) | **0.09**<br>**(0.05)** | — | **0.05**<br>**(0.03)** | 0.53<br>(0.03) | **0.15**<br>**(0.04)** | 0.05<br>(0.04) | **0.11**<br>**(0.05)** | — | **0.06**<br>**(0.03)** | **0.62**<br>**(0.04)** | 0.00 |
| horse ID + additive genetic + social group | 0.81 | **0.13**<br>**(0.03)** | 0.05<br>(0.03) | — | — | **0.09**<br>**(0.02)** | 0.53<br>(0.03) | **0.16**<br>**(0.04)** | 0.06<br>(0.04) | — | — | **0.12**<br>**(0.02)** | **0.66**<br>**(0.03)** | 4.86 |
| horse ID + additive genetic + social community | 0.89 | **0.12**<br>**(0.03)** | 0.04<br>(0.03) | **0.17**<br>**(0.04)** | — | — | 0.55<br>(0.02) | **0.13**<br>**(0.04)** | 0.05<br>(0.04) | **0.20**<br>**(0.04)** | — | — | **0.62**<br>**(0.04)** | 2.02 |
| horse ID + additive genetic | 0.81 | **0.12**<br>**(0.03)** | 0.07<br>(0.03) | — | — | — | 0.62<br>(0.02) | **0.15**<br>**(0.04)** | **0.09**<br>**(0.04)** | — | — | — | **0.76**<br>**(0.03)** | 31.03 |
| horse ID | 0.81 | **0.19**<br>**(0.02)** | — | — | — | — | 0.62<br>(0.02) | **0.24**<br>**(0.03)** | — | — | — | — | **0.76**<br>**(0.03)** | 33.58 |

PE: permanent environment, A: additive genetic, SC: social community, SP: spatial, SG: social group

Likelihood ratio tests of bivariate animal models in which we estimated additive genetic, permanent environment, and social community effects indicated phenotypic correlation between microbiota and gene family CAP traits (r_P_ = 0.71) was attributable to positive additive genetic (r_A_ = 0.98 ± 0.09 SE, χ^2^ = 4.27, p = 0.04; Table S5), permanent environment (r_sc_ = 0.99 ± 0.06 SE, χ^2^ = 16.60, *p* = 4.6*e*^−5^), social community (r_sc_ = 0.88 ± 0.04 SE, χ^2^ = 47.26, *p* = 6.2*e*^−12^), and residual correlations between traits (r_r_ = 0.54 ± 0.02 SE). Social correlation between traits was significantly <1, based upon likelihood ratio tests in which r_sc_ was constrained to 0.99 (χ^2^ = 8.89, *p* = 0.003).

We next partitioned variance in microbiota or gene family traits (CLR-transformed abundances), using models containing random effects for horse identity, social community similarity, and additive genetic relatedness (Figure 3). This model was selected as it outperformed more complex models for both microbiota and gene family CAP traits. Use of the same model structure also allowed for more direct comparisons between microbiome trait types. Permanent environment effects accounted for the lowest proportion of variance in microbe CLR-transformed abundances (mean *pe*^2^ = 0.04 ± 0.04 S.D.; maximum *pe*^2^ = 0.16 ± 0.04 S.E., *Sharpea azabuensis*), followed by additive genetic effects (mean *h*^2^ = 0.05 ± 0.05 S.D.; maximum *h*^2^ = 0.27 ± 0.05 S.E., Ca. *Campylobacter infans*), and social community effects (mean *sc*^2^ = 0.14 ± 0.07 S.D.; maximum *sc*^2^ = 0.59 ± 0.03 S.E., unclassified Basidiomycota fungi of the genus *Microbotryum*). Permanent environment effects likewise accounted for the lowest proportion of variance in gene family CLR abundance (mean *pe*^2^ = 0.02 ± 0.02 S.D.; maximum *pe*^2^ = 0.13 ± 0.04 S.E., alanine dehydrogenase), followed by additive genetic effects (mean *h*^2^ = 0.04 ± 0.04 S.D.; maximum *h*^2^ = 0.22 ± 0.04 S.E., anthranilate synthase), and social community effects (mean *sc*^2^ = 0.07 ± 0.05 S.D.; maximum *sc*^2^ = 0.23 ± 0.04 S.E., Dihydromethanophenazine:CoB— CoM heterodisulfide reductase). Significant heritability estimates after Benjamini-Hochberg FDR correction were observed for 563 of 1574 microbiota (36%) and 417 of 1416 gene families (29%) tested. Significant effects of social community were observed for 1443 of 1574 microbiota (92%; Appendix 3) and 896 of 1416 gene families tested (63%; Appendix 4). Significant maternal effects were not observed for any microbes or gene families tested (Appendix 5 & 6).

**Figure 3:**
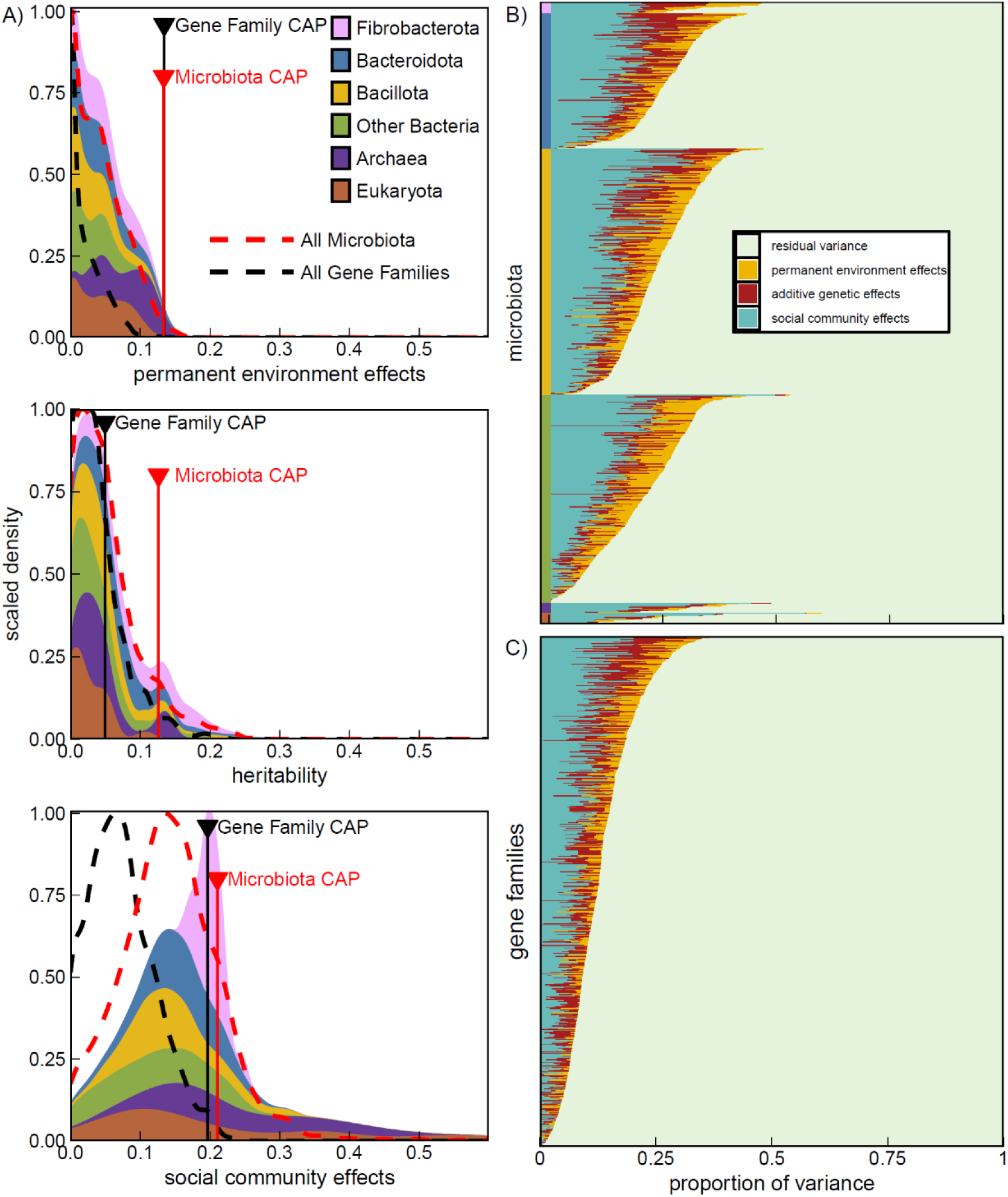
(A) density plots of proportional permanent environment, additive genetic, and social community contributions to variances in microbe centred log ratio (CLR) transformed abundance. Stacked density plots are coloured by major taxonomic grouping and scaled to the same total area per taxon. Dotted lines denote distribution of all microbes (red) or gene family (black) variance component estimates. Solid vertical lines capped with triangles denote the corresponding value estimate for survival-associated CAP axis microbiomes traits. Additionally, stacked barplots display the proportional effects of additive genetic, permanent environment, and horse social community effects on variances in CLR transformed abundances of (B) microbes or (C) gene families represented by different rows. Microbes are sorted according to major taxonomic grouping and coloured along the y-axis according to the legend in panel “A”. Variance component estimates were conditional on fixed effects for longitude (2^nd^ order polynomial), horse age (2^nd^ order polynomial), day of year (2^nd^ order polynomial), DNA extraction/library preparation plate (factor), and year of sample collection (factor).

Microbes and gene families more strongly associated with horse survival tended to be more strongly socially structured and repeatable within individuals (noting that repeatability arises from the combination of additive genetic and permanent environment differences among individuals). Specifically, we observed a negative relationship between the proportion of residual variance from animal models and the magnitude of the relationship between microbes or gene families and horse survival (microbes: β = −0.43 ± 0.03, t = −14.65, p = 2.0e^−16^, Figure 4; gene families: β = −0.12 ± 0.02, t = −5.67, p = 1.69e^−8^; Figure S4). The same patterns of non-independence were separately observed with respect to permanent environment effects (microbes: β = 0.12 ± 0.01, t = 9.19, p = 2.0e^−16^; gene families: β = 0.10 ± 0.01, t = 9.00, p = 2.0e^−16^), social community effects (microbes: β = 0.21 ± 0.02, t = 8.90, p = 2.0e^−16^; gene families: β = 0.17 ± 0.02, t = 8.06, p = 1.6e^−15^) and additive genetic effects for microbes (β = 0.09 ± 0.02, t = 5.31, p = 1.3e^−7^) but not gene families (β = 0.02 ± 0.02, t = 0.96, p = 0.34). These results were not attributable to differences in the relative abundance of microbiome traits, which was included in models as a covariate (Figure 4), since low abundance microbes or gene families may be less precisely estimated. As predicted, we observed that the proportion of unexplained phenotypic variance from animal models were strongly negatively associated with average log-transformed relative abundance among microbes (β = −0.03 ± 0.001, t = −22.16, p = 2.0e^−16^; Figure S5A). Contrary to expectations, we observed a positive parabolic relationship between residual variance proportions and average log transformed relative abundance of gene families (linear term: β = −0.35 ± 0.07, t = −5.24, p = 1.9e^−7^; quadratic term: β = 0.83 ± 0.06, t = 12.89, p = 2.0e^−16^), such that the lowest proportions of residual variance were observed among moderately abundant gene families (Figure S5B).

**Figure 4:**
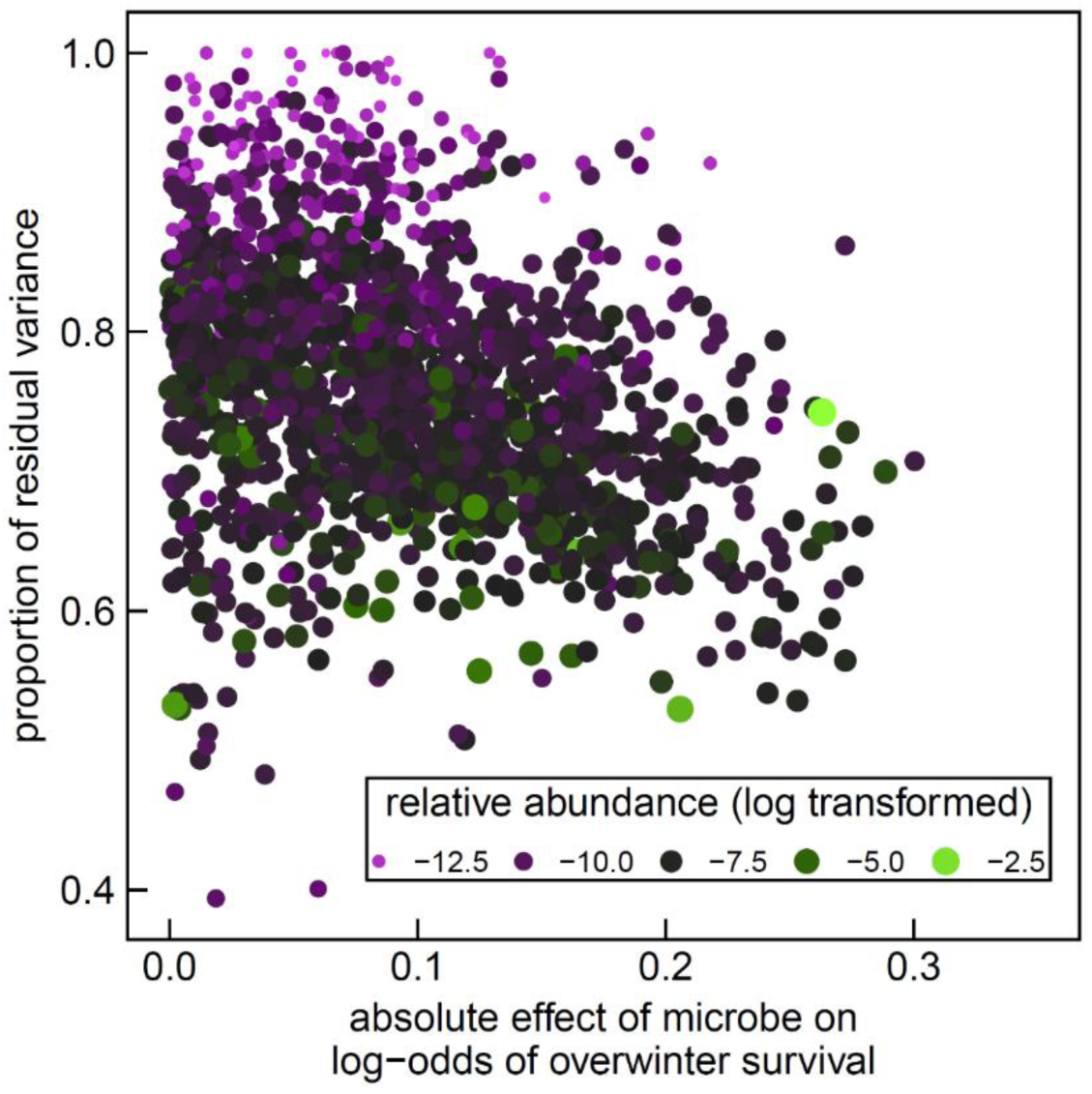
Scatterplot of the absolute magnitude of microbe associations with the odds of horse survival versus the proportion of residual variance remaining from animal models which contained terms for horse identity, pedigree-estimated genetic relatedness, and social community similarity, and fixed effects for longitude (2^nd^ order polynomial), horse age (2^nd^ order polynomial), day of year (2^nd^ order polynomial), DNA extraction/library preparation plate (factor), and year of sample collection (factor). Each point is colour and sized by its log-transformed relative abundance and represents the finest taxonomic bins to which reads could be classified.

## Discussion

We quantified patterns of inheritance for gut microbiome traits linked to fitness proxies in Sable Island feral horses. Microbiome gene family content was more strongly associated with fitness proxies than a microbiota derived CAP trait, supporting the ITSNTS hypothesis prediction that microbiome function is a more direct target of selection than microbiota membership^33^. We further demonstrate that fitness-related microbiome dimensions have a significant additive genetic basis, but contrary to our predictions and ecosystem-on-a-leash expectations^34^, we observed stronger evidence of narrow-sense heritability for microbiota community (h^2^ = 0.13 ± 0.04 S.E.) than gene family CAP traits (h^2^ = 0.05 ± 0.04 S.E.). In fact, significant narrow-sense heritability of gene family CAP trait values was only observed in the absence social community effects, or among more statistically powerful bivariate animal models. Therefore, the gene family CAP trait has an additive genetic basis, but its heritability exists near the detection limit for our dataset. Strong additive genetic covariance between CAP traits within a bivariate animal model suggests that same additive genetic basis underlies the heritability of fitness-associated microbiota and gene families.

Survival-associated microbiome CAP scores were more strongly heritable than the average microbe (mean *h^2^* = 0.05) or gene family trait (mean *h^2^* = 0.04). Average microbiome trait narrow-sense heritability in Sable Island horses is comparable to estimates from humans (mean *h^2^* = 0.05)^68^, domestic cattle (mean *h^2^* = 0.10)^69^ and wild yellow baboons (mean h^2^ = 0.06–0.08)^70^. Previous studies have not always accounted for social effects, which we found to contribute overestimation of microbiome trait heritability. Conversely, our observation-based pedigree may estimate genetic relationships with increased error when compared to genetic-based methods, thereby causing us to underestimate microbiome trait heritability^48^. Low average narrow-sense heritability of gut microbes has been used to conclude that host genetic variation is a minor contributor to overall microbiome variation within host populations^18^. However, we observed that fitness-associated microbiome traits had narrow-sense heritability estimates comparable to conventional vertebrate traits (morphology, physiology, phenology, behaviour)^71,72^, and so, should be capable of evolution in response to selection at the host level.

Social community structure was generally as important as combined additive genetic and permanent environment effects in determining fitness-associated microbiome trait variation. Properly accounting for social effects is clearly necessary to obtain accurate estimates of microbiome heritability, but our findings also highlight the need to consider mechanisms of non-genetic inheritance of microbiome trait variation (e.g., host-to-host microbe transmission)^73^. Social transmission of microbes is now recognized as an important determinant of microbiome structure in human and wild animal populations^21,24,27,29^. Although the social community effects we describe likely do not solely reflect microbiota transmission^74^, animal models containing a *social community* term outcompeted models containing more direct measures of shared environment (e.g., *spatial* or *social group identity* effects). The interpretation that the *social community* effects are related to microbe transmission is also consistent with previous eco-phylogenetic null modelling analyses, which identified bacterial dispersal limitation as a primary cause of microbiome dissimilarity between Sable Island horses^26^. Interestingly, we observed significant evidence for social structuring of both microbiota community and microbiome gene family traits. Bivariate analyses further identified high social covariance between survival-associated microbiota and gene family CAP trait measures (r_soc_ = 0.88), but the social correlation was significantly less than 1. This suggests that social relationships can influence both microbiome membership and functional potential^31^, but fitness-associated microbiota and gene family variation may not necessarily be transmitted through the same social networks (although they do socially covary). Overall, our results support theorization that metacommunity ecology understanding is necessary to predict host-microbiome evolution at micro-evolutionary scales^20,32^.

Permanent environment effects are important for natural selection but are not typically considered relevant to patterns of trait inheritance. However, permanent environment effects may be important for the inheritance of socially transmissible traits, such as traits derived from the microbiome^31^. Afterall, both genetic and permanent environment effects shape the repeatable microbial contributions that hosts make to their social network. Permanent environment effects were comparable in magnitude to additive genetic effects in shaping microbiota community CAP trait variation. In contrast, permanent environment effects appeared more important than additive genetic effects in determining gene family CAP trait variation. Priority effects and stabilizing ecological interactions between microbes likely both contribute to observations of permanent environment effects on microbiome traits^17,75,76^. Permanent environment effects may be caused by differences in development or early-life environments that lead to fixed physiological differences between individuals^77^. In some instances, microbes encountered in early life can themselves underlie permanent environment effects by programming host physiology in ways that persist into adulthood^78,79^. Maternal effects are another prominent determinant of the early life environment and one which commonly contributes to trait variation among animal populations^71^.

We observed no evidence for maternal identity effects on the Sable Island horse microbiome despite a strong biological expectation for maternal contributions to the gut microbiome^80–82^. Importantly, these results do not indicate that maternal effects do not contribute to the Sable Island horse microbiome; rather, they suggest that the contributions that mares make to their offspring’s adult microbiome may not be repeatable across births^83^. Weak evidence for maternal identity effects on microbiome traits has similarly been reported in wild yellow baboons^70^. It is possible that maternal identity effects on the gut microbiome may be diluted in gregarious species which have large social networks which, alongside mothers, shape the early life environment. However, even in non-gregarious wild wood mice (*Apodemus sylvaticus*) maternal effects on the microbiome in early life weaken through ontogeny^84^. Similar sensitivity of the horse microbiome to social relationships in the year of sample collection (rather than maternal identity or past social relationships) is evidenced by the performance of animal models containing *social community* effects which reflect turnover in social community membership across years.

Microbiota and gene family traits that were strongly related to horse survival also tended to be more strongly influenced by additive genetic, social community, and permanent environment effects (Figure 4; Figure S3). To assess the microbiome’s capacity to evolve in response to selection, previous studies have tended to focus on the average heritability of the microbes observed in the gut^16,18,70^. However, the heritability of the average microbe or microbial gene family may be less important for our understanding of adaptive evolution in the host-microbiome relationship than the specific heritability or social transmissibility of the microbiome traits most strongly connected to host fitness. Microbiome studies commonly seek to explain extrinsic and intrinsic causes of total microbiome variation contained within a population^85^. Yet, we found that the dimensions of the gut microbiome most strongly connected to host fitness components represented a small fraction of total microbiome variation. Rather than seeking to explain gut microbiome variation in its entirety, focusing on dimensions of the microbiome connected to host fitness or phenotypic variation may be more relevant to our understanding of microbiome-mediated adaptation in nature^86^.

Our observation of non-independence between fitness relationships and animal model variance component estimates likely has both a technical and biological basis. From a technical perspective, low abundance microbiome traits will be less precisely estimated^87^, causing increased error in both quantitative genetic animal models and selection analyses. Nonetheless, microbiota or gene family trait abundance did not explain relationships between animal model and selection analysis results. Non-independence between estimates of selection and inheritance could be partly an artefact of our sampling time scales^88^. Heritability and selection over annual periods (or lifetimes) can be most easily detected when acting on host traits that are highly repeatable over similar timescales^89^. Although selection may also act on mutable microbiome traits (e.g., rapid onset of gut dysbiosis)^90^, detecting such effects would require more frequent sampling across a smaller window of selection (e.g., survival across weeks rather than years). Our annual sampling timescale and use of inter-annual survival and reproductive success as fitness proxies may be biased towards detecting selection acting on highly repeatable microbiome traits.

From a biological perspective, gut-adapted microbes may simultaneously be more repeatable and capable of influencing the microbiome, utilizing microbe-, diet-, or host-derived substrates, and interacting with host physiology in ways that are consequential to host fitness, when compared to transient microbes not adapted to the gut^16,91^. Simultaneously, the adaptation of microbes to their host organism may make them more sensitive to heritable and non-heritable variation in host physiology^92^ and may even obligate their transmission more firmly to host social networks^30^. Gut-adapted microbe sensitivity to, and capacity to exert influence on, host physiology may therefore provide a biological explanation for observed non-independence between estimates of selection and our animal model results.

Non-independence between host fitness-microbiota relationships and microbiota inheritance patterns may be widespread across host-microbiome symbioses. For example, microbes that facilitate or inhibit methanogenesis are frequently reported as among the most highly heritable microbes within mammal populations^69,70,93–95^ and methanogenesis is known to be a key determinant of digestible energy flow in the animal gut^96,97^. We previously identified methanogenesis as a primary putative mechanism linking the gut microbiome to horse survival^15^, and now find that these same methanogenic features are among the most highly heritable and strongly socially structured microbiome traits within Sable Island horses. Although we observed evidence for directional selection against traits related to methane production during our seven-year study period, non-methane hydrogen sinks have physiological trade-offs (e.g., short-chain fatty acid profile changes, detrimental byproduct synthesis)^98^. The widespread heritability of microbiome traits related to methanogenesis could indicate that additive genetic variance in these traits is commonly maintained within mammalian populations by diversifying or fluctuating directional selection^99,100^.

In summary, we find evidence that natural selection acts more strongly on microbiome functional potential than community composition, but fitness-associated microbiota community variation is more faithfully transmitted across generations. Dimensions of the gut microbiome connected to horse survival and reproductive success are heritable among Sable Island horses, and so, theoretically capable of adaptive evolution in response to host-level selection. However, social effects were twice as important as additive genetic effects in determining fitness-associated dimensions of the microbiome. These results emphasize the need to consider the direct transmission of microbiota through social networks as an important non-genetic inheritance mechanism integral to shaping patterns of host-microbiome evolution in nature.

## Data availability

The shotgun metagenomic sequence that support the findings of this study have been deposited in the NCBI SRA under the BioProject accession codes PRJNA1102860 and PRJNA880353.

## Acknowledgements

We thank the students, research assistants, and volunteers who have contributed to the Sable Island Horse Project. We also thank the University of Calgary Center for Health Genomics and Informatics for assistance with library preparation and sequencing, as well as the Department of Fisheries and Oceans Canada (DFO), Canada Coast Guard, the Bedford Institute of Oceanography (DFO Science), Environment Canada, Parks Canada Agency, Maritime Air Charters Limited (Sable Aviation), and Sable Island Station (Meteorological Service of Canada) which provided in-kind and logistical support.

