## Supplementary Materials for "Host genetics and social relationships jointly shape fitness-associated microbiome variation in a population of feral horses"

Table S1: Results from a logistic regression mixed model to test the effects of faecal microbiota and non-microbiome predictors on the log odds of Sable Island horse overwinter survival. Models included a random effect for year of sample collection.

| | β | S.E. | z value | p value |
|---|---|---|---|---|
| intercept | 1.97 | 0.25 | 8.039 | $9.1e^{-16}$ |
| $CAP_{microbiota}$ | 0.62 | 0.07 | 9.228 | $2.0e^{-16}$ |
| sex (male) | 0.34 | 0.15 | 2.285 | 0.02 |
| mean longitude (linear term) | −5.87 | 2.93 | −2.001 | 0.05 |
| mean longitude (quadratic term) | −6.22 | 2.96 | −2.100 | 0.04 |
| age (linear term) | −14.71 | 3.24 | −4.544 | $5.5e^{-6}$ |
| age (quadratic term) | −15.54 | 2.97 | −5.237 | $1.6e^{-7}$ |
| parental status (male) | −0.59 | 0.19 | −3.202 | $1.0e^{-3}$ |
| parental status (female) | 0.13 | 0.26 | 0.477 | 0.63 |

Table S2: Results from a logistic regression mixed model to test the effects of faecal microbiome gene family and non-microbiome predictors on the log odds of Sable Island horse overwinter survival. Models included a random effect for year of sample collection.

| | β | S.E. | z value | p value |
|---|---|---|---|---|
| intercept | 2.05 | 0.26 | 8.016 | $1.1e^{-15}$ |
| $CAP_{gene\ family}$ | 0.86 | 0.07 | 11.569 | $2.0e^{-16}$ |
| sex (male) | 0.38 | 0.16 | 2.441 | 0.01 |
| mean longitude (linear term) | −6.82 | 2.99 | −2.280 | 0.02 |
| mean longitude (quadratic term) | −6.42 | 2.99 | −2.152 | 0.03 |
| age (linear term) | −14.33 | 3.32 | −4.311 | $1.6\ e^{-5}$ |
| age (quadratic term) | −17.05 | 3.05 | −5.596 | $2.2\ e^{-8}$ |
| parental status (male) | −0.67 | 0.19 | −3.539 | $4.0e^{-4}$ |
| parental status (female) | −0.02 | 0.27 | −0.061 | 0.95 |

Table S3: Results from a logistic regression mixed model to test the effects of faecal microbiota and non-microbiome predictors on the log odds of annual sex-specific reproductive success among Sable Island horses. Models included a random effect for year of sample collection.

| | $\beta$ | S.E. | z value | p value |
|---|---|---|---|---|
| intercept | −0.83 | 0.12 | −7.009 | $2.4e^{-12}$ |
| $CAP_{microbiota}$ : female | 0.34 | 0.08 | 4.236 | $2.3e^{-5}$ |
| $CAP_{microbiota}$ : male | 0.21 | 0.08 | 2.485 | 0.01 |
| sex (male) | −1.35 | 0.12 | −11.512 | $2.0e^{-16}$ |
| mean longitude (linear term) | −1.63 | 2.72 | −0.600 | 0.55 |
| mean longitude (quadratic term) | −4.28 | 2.80 | −1.528 | 0.13 |
| age (linear term) | 50.82 | 3.54 | 14.368 | $2.0e^{-16}$ |
| age (quadratic term) | −30.87 | 3.27 | −9.434 | $2.0e^{-16}$ |

Table S4: Results from a logistic regression mixed model to test the effects of faecal microbiome gene family and non-microbiome predictors on the log odds of annual sex-specific reproductive success among Sable Island horses. Models included a random effect for year of sample collection.

| | $\beta$ | S.E. | z value | p value |
|---|---|---|---|---|
| intercept | −0.87 | 0.12 | −7.380 | $1.59e^{-13}$ |
| $CAP_{gene\ family}$ : female | 0.49 | 0.08 | 6.005 | $1.91e^{-9}$ |
| $CAP_{gene\ family}$ : male | 0.15 | 0.08 | 1.816 | 0.07 |
| sex (male) | −1.29 | 0.11 | −11.240 | $2.0e^{-16}$ |
| mean longitude (linear term) | −2.33 | 2.73 | −0.854 | 0.39 |
| mean longitude (quadratic term) | −4.85 | 2.78 | −1.743 | 0.08 |
| age (linear term) | 51.46 | 3.56 | 14.456 | $2.0e^{-16}$ |
| age (quadratic term) | −31.75 | 3.28 | −9.704 | $2.0e^{-16}$ |

Table S5: Variance component estimates from a bivariate animal model of microbiota and gene family CAP traits conditioned on fixed effects for longitude (2nd order polynomial), horse age (2nd order polynomial), days since July 1st of sampling year (2nd order polynomial), DNA extraction/library preparation plate (factor), and year of sample collection (factor). Raw variance component estimates and proportional contribution towards to phenotypic variance reported with standard error in brackets.

<table>
<tr><th>Trait</th><th>$V_A$</th><th>$V_{PE}$</th><th>$V_{SC}$</th><th>$V_R$</th><th>$COV_A$</th><th>$COV_{PE}$</th><th>$COV_{SC}$</th><th>$COV_R$</th><th>$h^2$</th><th>$pe^2$</th><th>$sc^2$</th><th>$r^2$</th></tr>
<tr><td>$CAP_{microbiota}$</td><td>$5.9e^{-4}$ $(2.5e^{-4})$</td><td>$9.0e^{-4}$ $(2.5e^{-4})$</td><td>$1.6e^{-3}$ $(2.8e^{-4})$</td><td>$2.9e^{-3}$ $(1.4e^{-4})$</td><td rowspan="2">$3.7e^{-4}$ $(1.9e^{-4})$</td><td rowspan="2">$7.6e^{-4}$ $(1.9e^{-4})$</td><td rowspan="2">$1.1e^{-3}$ $(2.1e^{-4})$</td><td rowspan="2">$1.5e^{-3}$ $(1.1e^{-4})$</td><td>0.10 (0.04)</td><td>0.15 (0.04)</td><td>0.27 (0.04)</td><td>0.48 (0.03)</td></tr>
<tr><td>$CAP_{gene\ family}$</td><td>$2.5e^{-4}$ $(1.6e^{-4})$</td><td>$6.4e^{-4}$ $(1.8e^{-4})$</td><td>$9.9e^{-4}$ $(2.0e^{-4})$</td><td>$2.8e^{-3}$ $(1.2e^{-4})$</td><td>0.05 (0.03)</td><td>0.14 (0.04)</td><td>0.21 (0.04)</td><td>0.60 (0.03)</td></tr>
</table>

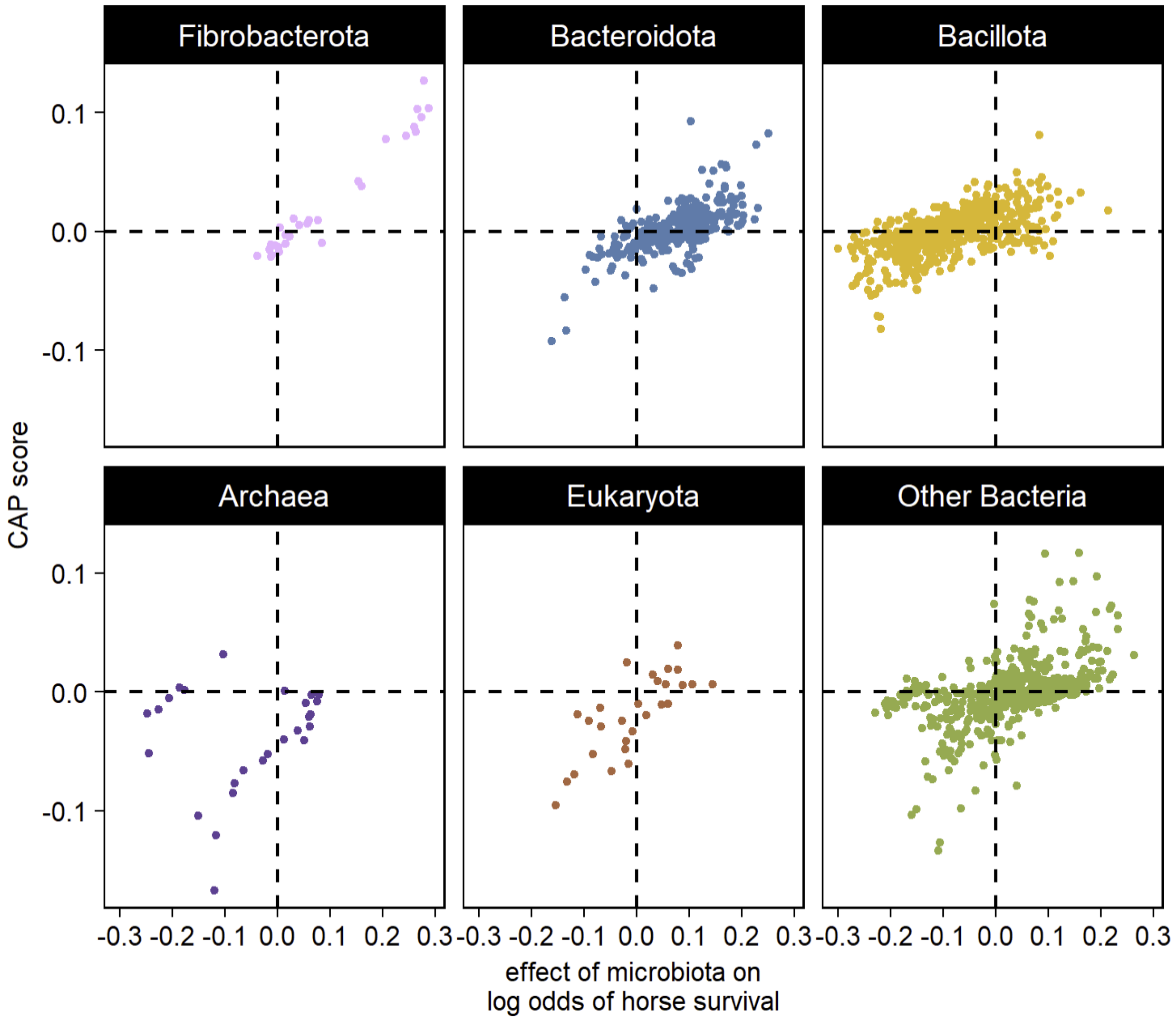


Figure S1: Microbe centred log ratio transformed abundance (centred and variance standardized) associations with the log-odds of horse survival versus microbe score along a survival constrained canonical analysis of principal coordinates (CAP) axis. Points denote taxonomic bins representing the finest levels to which reads could be classified, coloured by major taxonomic grouping.

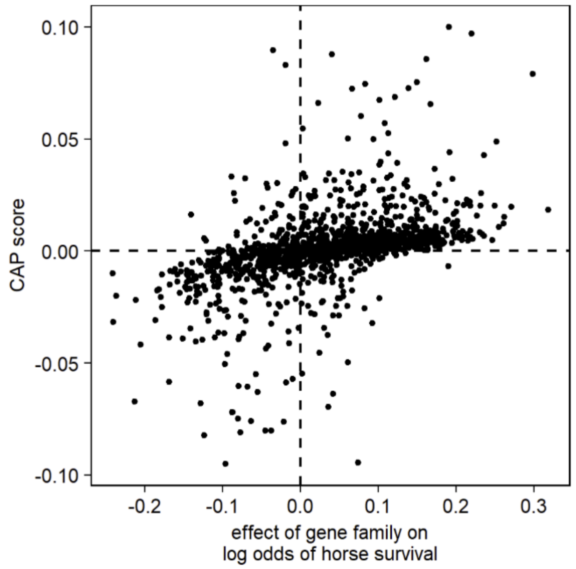


Figure S2: Gene family centred log ratio transformed abundance (centred and variance standardized) associations with the log-odds of horse survival versus gene family CAP score along a survival constrained canonical analysis of principal coordinates (CAP) axis.

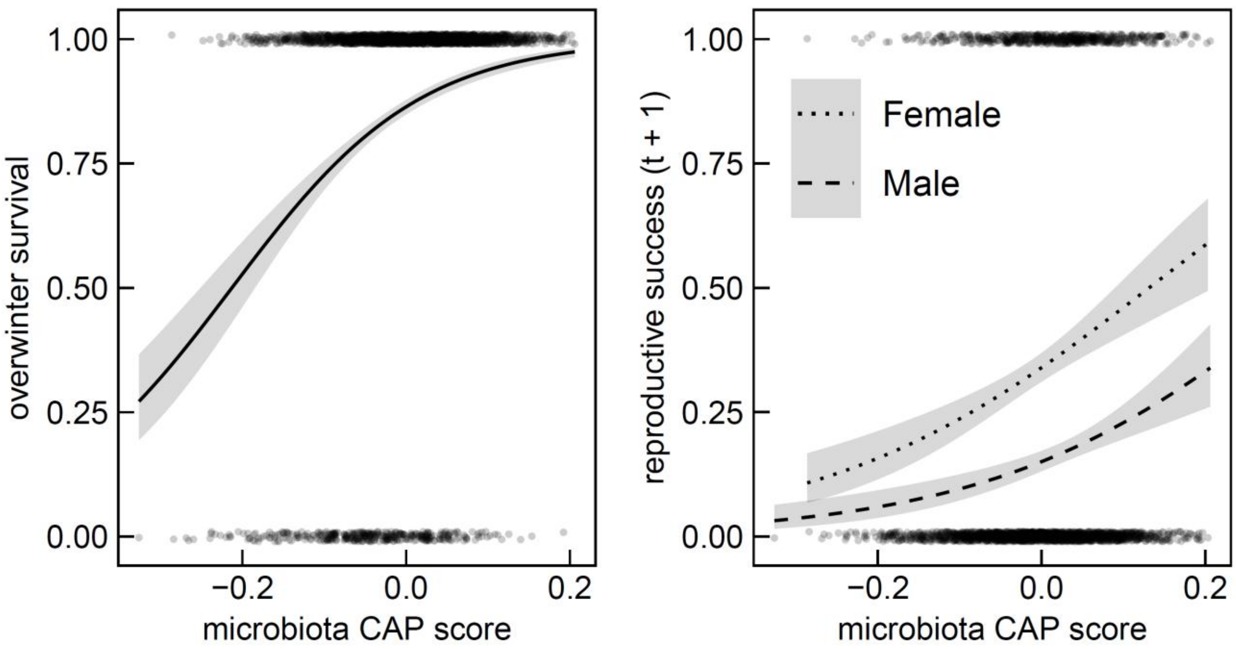


Figure S3: Jittered scatterplots of (a) horse overwinter survival and (b) sex-specific reproductive success in the year following faecal sample collection in response to weighted average scores of samples from a canonical analysis of principal coordinates (CAP) of microbiota profiles constrained by overwinter survival. Points denote individual samples with best-fit logistic regression lines and 95% confidence interval shading.

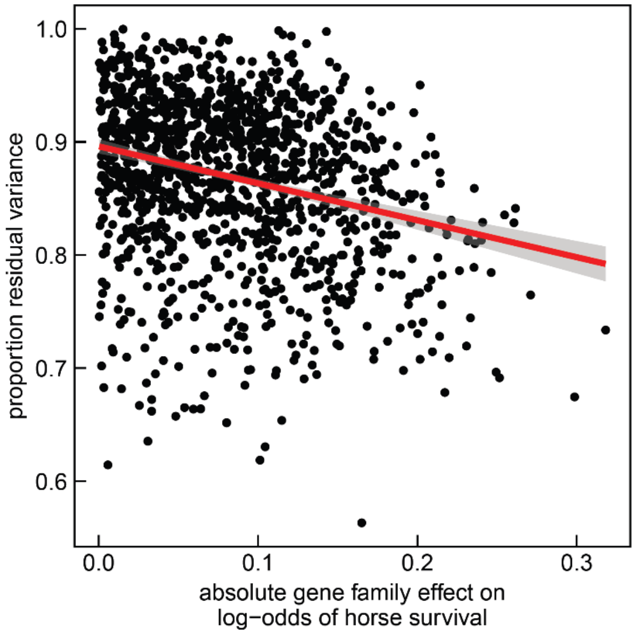


Figure S4: Scatterplot of the absolute magnitude of gene family associations with the odds of horse survival versus the proportion of residual variance remaining from animal models which contained random effects for horse identity, pedigree-estimated genetic relatedness, and social community similarity, and fixed effects for longitude (2nd order polynomial), horse age (2nd order polynomial), day of year (2nd order polynomial), DNA extraction/library preparation plate (factor), and year of sample collection (factor). Each point represents a different gene family. The red line denotes the best fit linear relationship with 95% confidence interval shading.

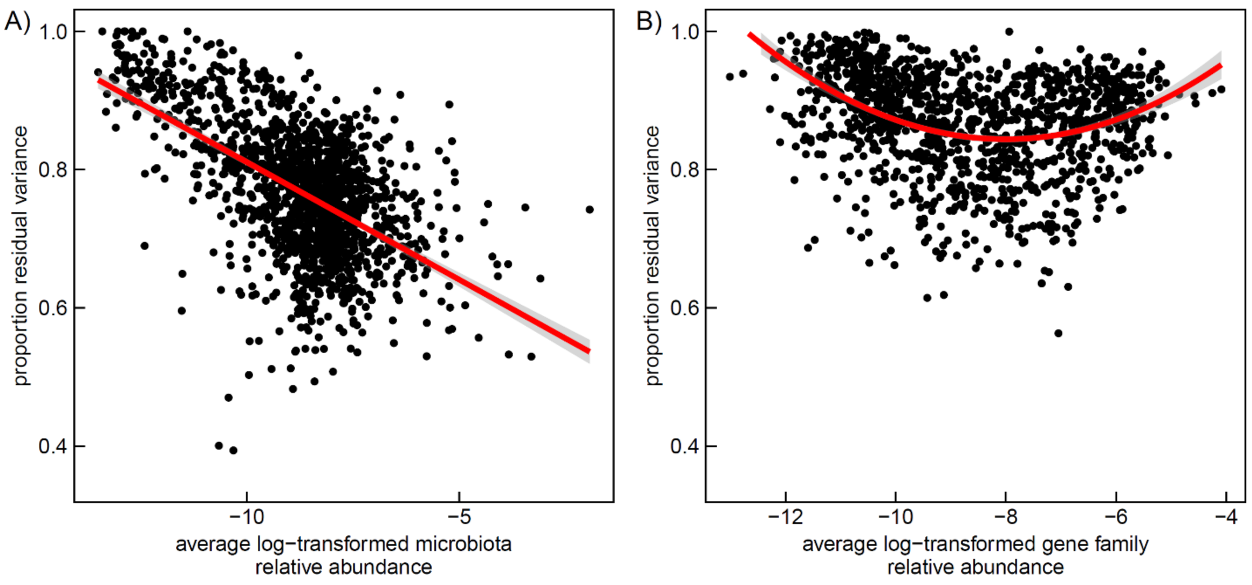


Figure S5: Scatterplots of proportions of residual variance in (A) microbe or (B) gene family abundance from animal models (centred log ratio transformed) versus the average relative abundance of microbiome traits (log transformed). Solid red lines denote the best fit linear (panel A) and quadratic (panel B) lines with 95% confidence interval shading. Animal models were conditioned on fixed effects for longitude (2nd order polynomial), horse age (2nd order polynomial), day of year (2nd order polynomial), DNA extraction/library preparation plate (factor), and year of sample collection (factor), and random effects for random effects for horse identity, pedigree-estimated genetic relatedness, and social community similarity.
